# Vagus Nerve Stimulation Promotes Gastric Emptying by Increasing Pyloric Opening Measured with Magnetic Resonance Imaging

**DOI:** 10.1101/253674

**Authors:** Kun-Han Lu, Jiayue Cao, Steven Oleson, Matthew P Ward, Terry L Powley, Zhongming Liu

## Abstract

**Background:** Vagus nerve stimulation (VNS) is an emerging electroceutical therapy for remedying gastric disorders that are poorly managed by pharmacological treatments and/or dietary changes. Such therapy seems promising since the vagovagal neurocircuitry controlling the enteric nervous system strongly influences gastric functions.

**Methods:** Here, the modulatory effects of left cervical VNS on gastric emptying in rats was quantified using a 1) feeding protocol in which the animal voluntarily consumed a post-fast, gadolinium-labeled meal and 2) newly developed, robust, sensitive and non-invasive imaging strategy to measure antral motility, pyloric activity and gastric emptying based on contrast-enhanced magnetic resonance imaging (MRI) and computer-assisted image processing pipelines.

**Key Results:** VNS significantly accelerated gastric emptying (control vs. VNS: 24.9±3.5% vs. 40.7±3.9% of meal emptied per 4hrs, p<0.05). This effect resulted from a greater relaxation of the pyloric sphincter (control vs. VNS: 1.4±0.2 vs. 2.5±0.5 mm^2^ cross-sectional area of lumen, p<0.05), without notable changes in antral contraction amplitude (control vs. VNS: 30.6±3.0% vs. 32.5±3.0% occlusion), peristaltic velocity (control vs. VNS: 0.67±0.03 vs. 0.67±0.03 mm/s), or frequency (control vs. VNS: 6.3±0.1 vs. 6.4±0.2 cpm). The degree to which VNS relaxed the pylorus positively correlated with gastric emptying rate (r = 0.5465, p<0.01).

**Conclusions & Inferences:** The MRI protocol employed in this study is expected to enable advanced preclinical studies to understand stomach pathophysiology and its therapeutics. Results from this study suggest an electroceutical treatment approach for gastric emptying disorders using cervical VNS to control the degree of pyloric sphincter relaxation.

**Key Points:**

1. Vagus nerve stimulation is emerging as a new electroceutical therapy for treating gastric disorders. However, its underlying mechanism(s) and therapeutic effect(s) remain incompletely understood.
2. Vagus nerve stimulation significantly accelerated gastric emptying by promoting the relaxation of the pyloric sphincter.
3. MRI offers high spatial and temporal resolution to non-invasively characterize gastric motility and physiology in preclinical animal models.

## Introduction

Dysregulation of gastrointestinal (GI) function is associated with gastroparesis (1, 2), obesity (3) gastroesophageal reflux disease and other GI disorders (4). These diseases are often chronic, idiopathic, and to date, not readily mitigated by surgical, pharmaceutical, and/or dietary treatments (5). For this reason, researchers have begun to explore electrical stimulation of the vagus nerve as an alternative strategy to remedy gastric disorders (6-10).

The vagus nerve plays an important role in mediating the interaction between the central nervous system (CNS) and the gut. It carries coordinated afferent information from the GI tract to the nucleus tractus solitarius (NTS) in the brainstem, and issues efferent signals from the dorsal motor nucleus of the vagus (DMV), primarily to the lower third of the esophagus and the stomach to modulate their functions (11-14). Different branches of the vagus innervate different parts of the GI tract. Therefore, fiber-selective or feedback-controlled stimulation of the vagus is potentially a favorable and effective approach to treat targeted gastric disorders.

Substantial efforts have been made in mapping the topographical and functional organization of the vagus nerve (15, 16). Initial promise of VNS has also been demonstrated for treating epilepsy (17), anxiety disorders (18), chronic heart failure (19), apnea (20) and inflammation (21). However, the therapeutic potential of VNS on gastric disorders remains unclear and its mechanism-of-action is still elusive. The reasons are likely multiple, but two stand out: First, the GI tract is not only innervated by the extrinsic vagovagal neurocircuits, but also by its intrinsic enteric network. The interplay of extrinsic and intrinsic innervations are rather complex (11). Second, different gastric regions are innervated by different vagal branches, thereby having differential responses to VNS, yet proper functioning of the GI tract is thought to require a coordinated choreography of all gastric regions (15).

Most existing neuromodulation protocols measure physiological responses to stimulation at discrete locations without considering the global state of the stomach (22-24). In this regard, magnetic resonance imaging (MRI) yields simultaneous surveys of many tissues, making it an ideal tool to study various aspects of gastric motility. While gastric MRI has been applied to humans (25, 26), it has rarely been used in small animal models for preclinical assessment. We recently developed a contrast-enhanced animal MRI technique for comprehensive assessment of gastric anatomy, motility, emptying and nutrient absorption in small animals (27). Adopting this technique, we set out to evaluate the effects of cervical VNS on antral contraction, pyloric opening and gastric emptying in heathy rats. The proposed experimental protocol and findings could offer new insights in the use of animal gastric MRI to evaluate the efficacy of therapeutics in treating gastric disorders.

## Materials and Methods

Briefly, a gadolinium-based contrast agent was mixed with the animal’s meal in order for chyme to appear “bright” in MRI scans, thereby delineating the gastric and intestinal volume. A multi-slice MRI sequence was used to scan the GI volume with high spatial resolution, and a similar sequence with a smaller spatial coverage was used to scan antral contractions with high temporal resolution. Measurements of gastric functions and physiology included the overall change in GI volume, gastric emptying, forestomach volume, corpus volume, antral volume, antral contraction frequency, antral peristaltic wave velocity, antral contraction amplitude, pyloric opening size, intestinal filling and, indirectly, absorption.

### Animal Protocol

Twenty-one rats (Sprague-Dawley, male, 228-330g) were included in the study. All study procedures were approved by the Purdue Animal Care and Use Committee. Rats were housed individually in ventilated cages with elevated stainless steel wire floors to prevent the animals from accessing their feces. The environment was maintained on a 12:12h light-dark cycle (lights on at 6AM and lights off at 6PM).

### Animal Preparation and Surgical Protocol

According to a feeding protocol described elsewhere (27), each animal was trained until (approximately 7 days) it would voluntarily consume a fixed quantity (10g) of DietGel (DietGel Recovery, ClearH2O, ME, USA). On the day for gastric MRI, each animal was given the test meal with a mixture of dietgel and an MRI contrast agent – Gadolinium (Gd). Specifically, 10g dietgel was liquefied through double-boiling in warm water at 45°C and mixed with 22.4mg Gd-DTPA powder (#381667, Sigma Aldrich, St. Louis, USA). The liquefied dietgel solution was cooled to room temperature to return it to the semisolid gel state.

Following an over-night food restriction (18 hours, 5PM to 11AM), rats were able to voluntarily consume the Gd-labeled test meal. Then, each animal was anesthetized with 4% isoflurane mixed with oxygen at a flow rate of 500ml/min for 5 minutes. Of the 21 rats, 10 underwent neck surgery for implantation of a bipolar cuff electrode around the left cervical vagus nerve. After administering a preoperative bolus of carprofen (10mg kg^−1^, IP; Zoetis, NJ, USA) and performing a toe-pinch to assure adequate anesthesia, a ventral midline cervical incision was made between the mandible and sternum. The subcutaneous tissue was then bluntly dissected and retracted laterally together with the mandibular salivary glands to reveal the trachea and the left carotid artery. Upon exposure of the left carotid artery, the left cervical vagus nerve, which sits lateral and runs parallel to the carotid artery above the level of the carotid bifurcation, was identified. The connective tissues surrounding the left cervical vagus nerve were carefully dissected so that a 10-15 mm portion of the cervical vagal trunk was isolated from the carotid artery. A custom-designed bipolar cuff electrode (MicroProbes, Gaithersburg, MD, USA) with a platinum-iridium wire lead was wrapped and secured on the isolated vagus nerve. The lead was externalized prior to suturing the incision site.

The animal was then placed in prone position on a water-heated MR-compatible cradle. On the cradle, the animal received a bolus injection of 0.01mg kg^−1^ dexmedetomidine solution (0.05mg ml^-1^, SC; Zoetis, NJ, USA). Five minutes later the isoflurane dose was reduced to 0.3-0.5% isoflurane mixed with oxygen at a flow rate of 500ml/min. Fifteen minutes after the initial bolus, a continuous subcutaneous infusion of dexmedetomidine was administered (0.03mg kg^-1^ hour^−1^, SC). An MRI-compatible system (SA Instruments Inc., Stony Brook, NY, USA) was used to monitor respiration, cardiac pulsation, and body temperature to ensure a stable physiological state throughout the experiment. The two leads of the vagal electrodes were connected to a pair of twisted wires that ran from the MRI bore to the console room, and the wires were further connected to a constant-current stimulator (A-M Systems Model 2200). Upon the start of the first MRI acquisition, electrical pulses (monophasic pulses with alternating polarity, inter-pulse duration (IPD) = 50ms; pulse amplitude (PA) = 0.6mA; pulse width (PW) = 0.36ms; frequency = 10Hz; 20 seconds on and 40 seconds off) were delivered to the cervical vagus throughout the 4-hour experiment.

### Gastric MRI

The animals were scanned in a 7-tesla horizontal-bore small-animal MRI system (BioSpec 70/30; Bruker Instruments, Billerica, USA) equipped with a gradient insert (maximum gradient: 200mT m^-1^; maximum slew rate: 640T m^-1^ s^-1^) and a volume transmit/receive ^1^H RF coil (86 mm inner-diameter).

As in our earlier study (27), after the long axis of the stomach was localized with the initial MRI scans, the MRI protocol was performed with a series of alternating volumetric scans and fast scans; the former was for quantifying gastric volume with higher spatial resolution and larger spatial coverage, whereas the latter was for assessing antral motility with higher temporal resolution and more targeted spatial coverage. The volumetric scans were acquired using a two-dimensional Fast Low Angle Shot gradient echo (FLASH) sequence with repetition time (TR) = 124.131 ms, echo time (TE) = 1.364 ms, flip angle (FA) = 90°, 30 oblique slices, slice thickness = 1 mm, FOV = 60 × 60 mm^2^, in-plane resolution = 0.23 × 0.23 mm^2^, and 4 averages. The fast scans were acquired using a two-dimensional FLASH sequence with TR/TE = 11.784/1.09ms, FA = 25°, four oblique slices, slice thickness = 1.5mm, FOV = 60 × 60 mm^2^, in-plane resolution 0.47 × 0.47 mm^2^, no averaging, and 150 repetitions. The four fast scan slices were positioned and adjusted to cover the antrum, pylorus and duodenum, based on the immediately preceding volumetric images to account for the stomach displacement during gastric emptying.

To minimize motion artifacts, both volumetric and fast scans were respiration-gated such that images were acquired during end-expiratory periods, while the chest volume stayed roughly unchanged. With the respiratory gating, the volumetric scan took about 4 minutes; the fast scan took ∼2 seconds per repetition and lasted ∼6 min for 150 repetitions. The volumetric and fast scans were repeated in an interleaved manner for a total of four hours.

### Assessment of GI volume, compartmental volume, and emptying rate

The GI volume was assessed globally and compartmentally, which included the gastric volume and intestinal volume. The intestinal volume comprised the duodenal, jejunal, and ileal volumes. The gastric volume was further partitioned into forestomach, corpus and antral volumes. The volumes were sampled approximately every 15 minutes for 4 hours. Specifically, the contrast-enhanced lumenal volume of the GI tract at different times were segmented, partitioned, and quantified separately from the volumetric scans by using an image processing pipeline (27). The volume of each compartment measured at intervals was further normalized as a percent of its initial volume at time 0. This normalization step allowed us to observe the relative volume change over time for each animal, while accounting for the varying amount of the meal intake for different animals. The time series of gastric volumes were resampled at 15-min intervals for every animal and then averaged across animals to characterize gastric emptying at the group level.

### Assessment of gastric motility

The frequency, amplitude and velocity of the peristaltic wave in the gastric antrum were quantified from the fast scans by using a custom-built Matlab image processing pipeline (27). Briefly, the antrum was first delineated from a stack of 4 slices, and the proximal-to-distal antral axis was determined. The cross-sectional areas perpendicular to the proximal-to-distal antral axis were then calculated by summing the number of antral voxels within each cross-sectional plane. By iteratively doing so for each volume, we obtained a time series that represented the cross-sectional area (CSA) change of the antrum at different locations distant to the pylorus. In the CSA time series, the maxima of the time series indicated antral distension and the minima antral contraction (i.e. the lumen was largely occluded by the depth of the constriction of the antral wall). The antral contraction frequency, occlusion amplitude and velocity were computed from the time series. In this study, the antral motility indices were obtained from the middle antrum, which was 4.7mm distant from the pylorus. Of the 11 animals in the control group, 5 animals were excluded from the study due to either disoriented antrum or the presence of gastric air in the antrum. One animal in the VNS group was excluded from the study using the same exclusion criteria.

### Measurement of the size of the pyloric sphincter lumen

To measure the size of the pyloric sphincter, we manually determined a cross-sectional plane that was perpendicular to the outflow direction of the terminal antrum on the segmented GI tract from the volumetric scans. The CSA of the pyloric sphincter was calculated by counting the number of lumenal voxels in the determined plane, as shown in Fig. 3A. This process was repeated for each volume, and a time series that characterized the pyloric opening was obtained for each animal.

### Statistical analysis

Unless otherwise stated, all data are reported as mean±standard error of mean (SEM). A probability (p-value) < 0.05 was considered significant to reject the null hypothesis. To evaluate the significance of the difference in the gastric emptying profile between the two conditions, the emptying curve from each subject was modeled by a Weibull distribution expressed as below, for which the two parameters (*t*_const_, *β*) were estimated by the least-squares method (28),

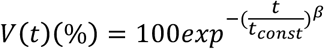

where V(t) is the remaining volume at experiment time *t* (min), *β* is the shape parameter of the curve, and *t*_const_ is the emptying time constant (min). For example, when *β* = 1, *t*_const_ represents the time estimated to empty ∼63% of the initial volume. The fitting was done by using the *fit* function in MATLAB. Only the estimated parameters with goodness of fit (R^2^) metrics greater than 0.85 were subject to the subsequent statistical analysis. A two-sample Student’s t-test was performed to assess the significance of the differences between the fitted parameters for the two conditions. To further assess the difference of remaining volume in each compartment, one-way ANOVA was conducted on individual time points between the two conditions. One-way ANOVA was also applied to determine whether there were statistically significant differences in antral motility indices and the degree of the pyloric opening between the control and the VNS conditions.

## Results

### Gd-labeled meal revealed the gastric lumen in MRI

All animals voluntarily consumed the Gd-labeled Dietgel (mean±SE: 7.72±0.39g; 6.08±0.31ml) in 13±6 minutes on the day of imaging following 7 days of training. Eleven control rats were scanned for gastric MRI under normal physiology, whereas the remaining 10 rats were scanned with their left cervical vagus nerve being electrically stimulated for 4 hours (Fig. 1A). Figure 1B shows the electrical stimulation paradigm. Adding gadolinium to the meal shortened the T1 relaxation. The meal thus appeared with much higher image intensity on the T1-weighted images compared to the surrounding tissues (Fig. 1C, 1D). The gastric volume and motility were quantified from our previously established image processing pipeline (27). Animals that underwent acute electrode implant surgeries prior to imaging were euthanized at the end of the experiment.

**Figure 1.**
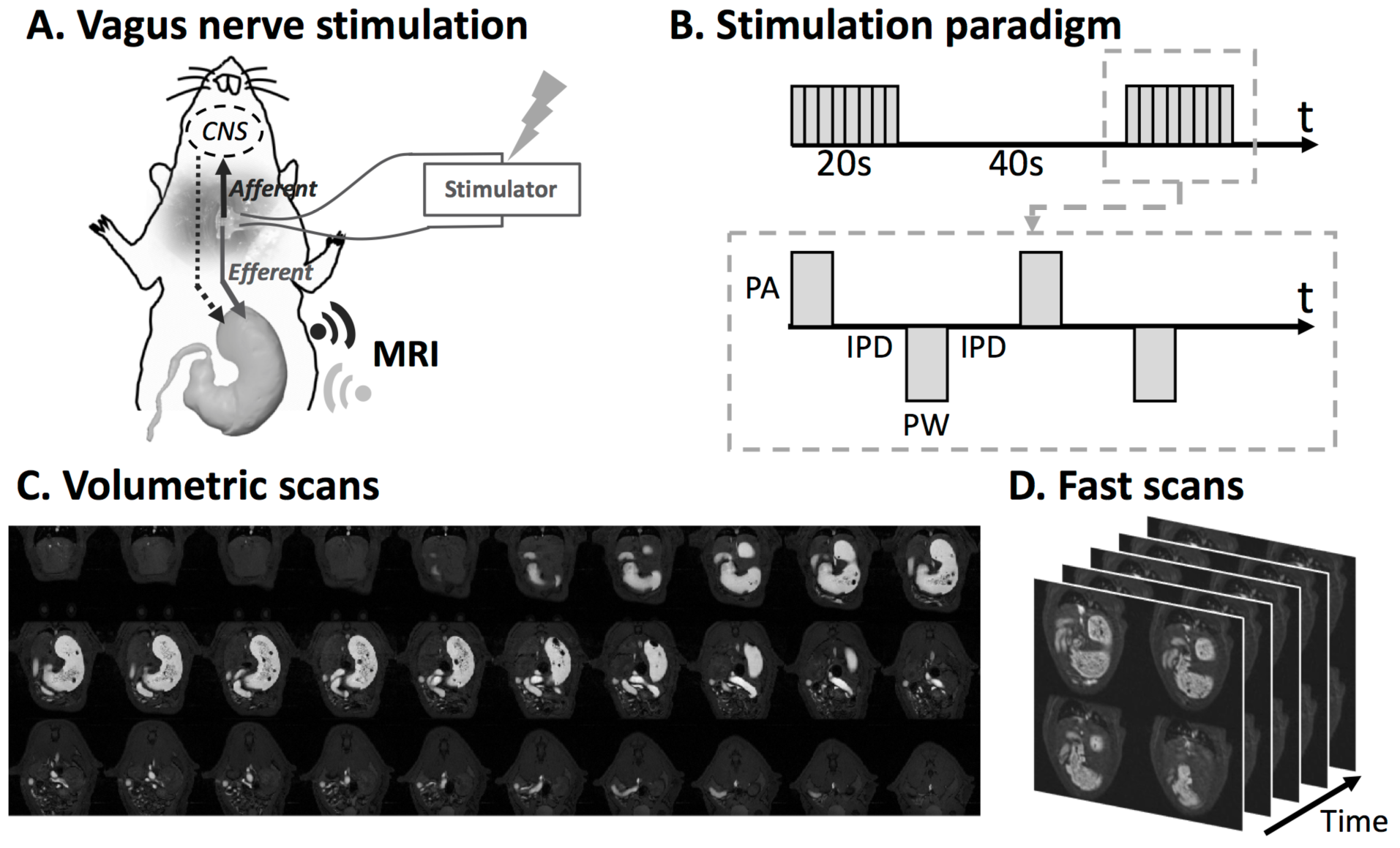
Vagus nerve stimulation and imaging sequence. **A.** Schematic diagram of the experiment protocol. A bipolar cuff electrode was implanted at the left cervical vagus of a rat. Vagus nerve stimulation was performed during the 4-hour MRI scans. **B.** Stimulation waveform, paradigm and parameters used in this study. (inter-pulse duration (IPD) = 50ms; pulse amplitude (PA) = 0.6mA; pulse width (PW) = 0.36ms; frequency = 10Hz; 20 seconds on and 40 seconds off) **C.** Example semi-coronal view images from the 2D multi-slice volumetric scan. The gastrointestinal tract is highlighted by the Gadolinium-labeled meal. **D.** Fast scans of the antrum with four slices. The position of the slice package was determined from images acquired from the volumetric scan.

### Vagus nerve stimulation promoted gastric emptying

Fig. 2 shows the modulatory effect of VNS on gastric emptying. All volumes were normalized against the volume within each compartment at time 0 – which indicates the emptying rate with respect to their initial volume – and then averaged across animals. Fig. 2A illustrates the 3D GI tract rendered from the high resolution volumetric scans. Different compartments of the GI tract were separated and labeled by the processing pipeline. Fig. 2B shows the volume change (%) of the GI tract (gastric volume + intestinal volume). The volume changes in the GI tract is strictly determined by the meal volume, secretion volume, and the volume absorbed by the intestines. In the first 30 minutes, there was an increase in the total GI volume in the control group (likely due to secretion) but not in the VNS group. After approximately one hour, the rate of volume change was faster in the VNS group than in the control group. Overall, gastric emptying was faster with VNS compared to the control condition (Fig. 2C). A compartment-wise analysis showed that the emptying rate in the forestomach (Fig. 2E), corpus (Fig. 2F), and antrum (Fig. 2G) was increased by VNS. Note that the antral volume stayed almost unchanged without VNS, suggesting a balance between the delivery of chyme to the antrum and the transfer of chyme to the duodenum. Such a balance was not observed in the VNS condition, where the trans-pyloric flow was faster than the antral emptying. In Fig. 2D, the intestinal volume (i.e., duodenum, jejunum and ileum) increased as the chyme filled the intestines, especially during the first 30 minutes. The intestinal volume stayed roughly unchanged from 60-240 minutes, reflecting a balance between intestinal filling and absorption. No significant difference in intestinal volume was observed between the control baseline and the VNS conditions.

**Figure 2.**
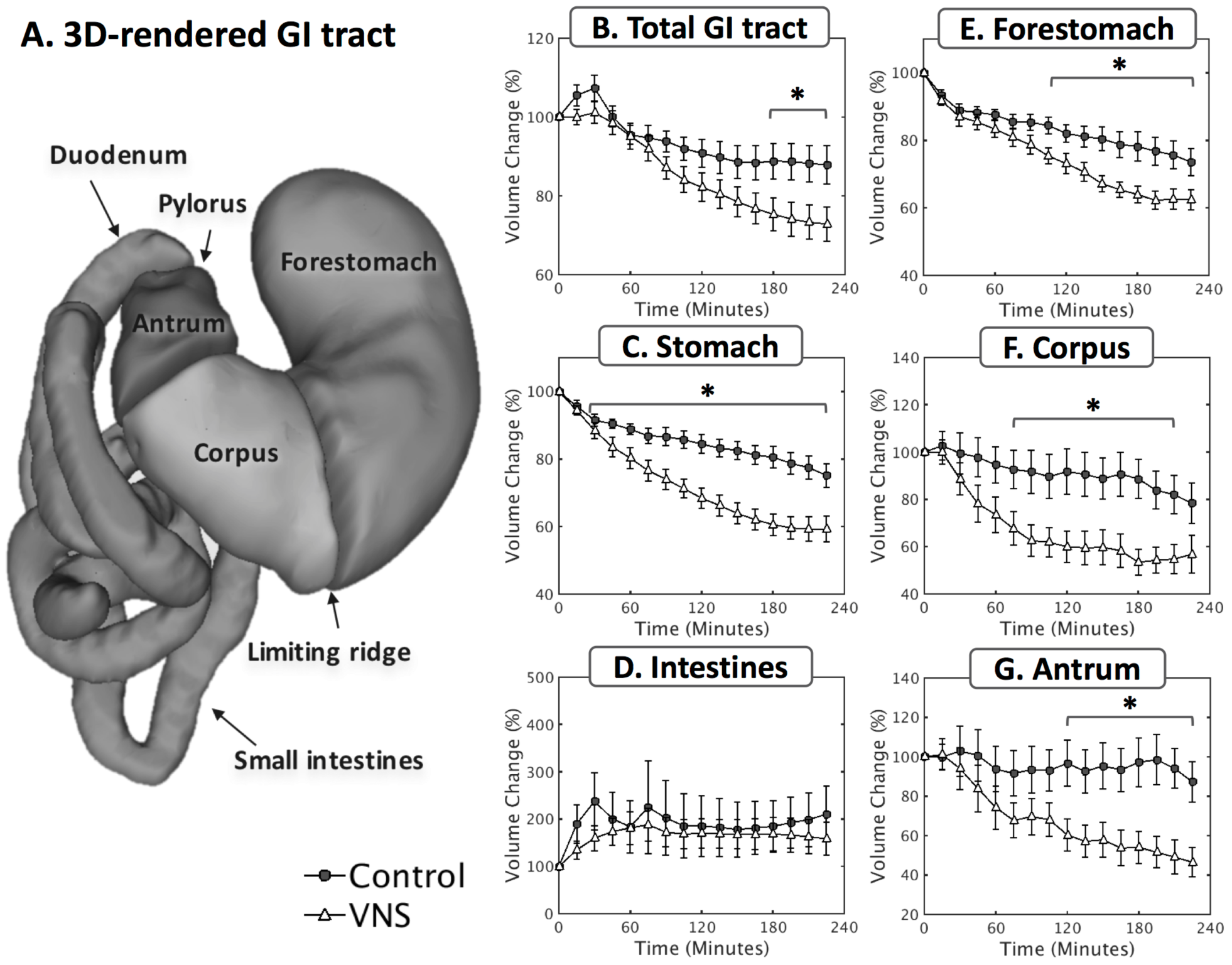
Vagal stimulation significantly enhances the rate of gastric emptying. **A.** 3D volume rendering of the lumen of the gastrointestinal tract. The profiles of volume change were quantified at a global scale including **B.** total GI tract, **C.** stomach, **D.** intestines, as well as at a compartment-wise scale including **E.** forestomach, **F.** corpus, **G.** antrum. *: p<0.05.

The gastric emptying curves were quantitatively characterized by a shape constant (*β*) and a time constant (*t*_const_). No significant difference in shape constant was found between the two conditions [control (n=8) vs. VNS (n=9): 1.04±0.08 vs. 0.85±0.03]. The time constant was significantly less in the VNS condition than in the control condition [control (n=8) vs. VNS (n=9): 882±56 vs. 452±15 min, p<0.05], suggesting that VNS increased the rate of gastric emptying. The difference of volume change within each compartment were also statistically evaluated as illustrated in Fig.2B-G (*p<0.05). There was a significant difference in the overall volume change in the GI tract (control vs. VNS: 12.1±4.8% vs. 27.2±4.3%, p<0.05), in the stomach (control vs. VNS: 24.9±3.5% vs. 40.7±3.9%, p<0.05), in the forestomach (control vs. VNS: 26.4±4.1% vs. 37.6±3.0%, p<0.05), in the corpus (control vs. VNS: 21.7±8.5% vs. 43.2±7.9%, p<0.05), and in the antrum (control vs. VNS: 12.7±10.2% vs. 53.5±7.3%, p<0.05) between the two conditions.

### Vagus nerve stimulation increased the size of the pyloric lumen

The lumenal CSA of the pyloric ring was defined as shown in Fig. 3A. Figure 3B shows how the size of the pylorus size changed during the 4-hour experiment for VNS animals and their controls. With VNS, the lumenal CSA of the pylorus was on average significantly greater than that without VNS (control vs. VNS: 1.4±0.2 vs. 2.5±0.5 mm^2^; p<0.05; Fig. 3C), particularly at t = 15, 75, 90, 135, 150 and 180 minute (p<0.05, uncorrected). The increase in the pyloric lumenal size was further correlated with the increase in stomach emptying. The overall volume change (%; four-hour difference) in the stomach was significantly correlated with the CSA of the pyloric lumen (r = 0.5465, p < 0.01). Note that VNS in general lead to a larger volume change in the stomach and to a greater size of the pylorus as shown in Fig. 3D.

**Figure 3.**
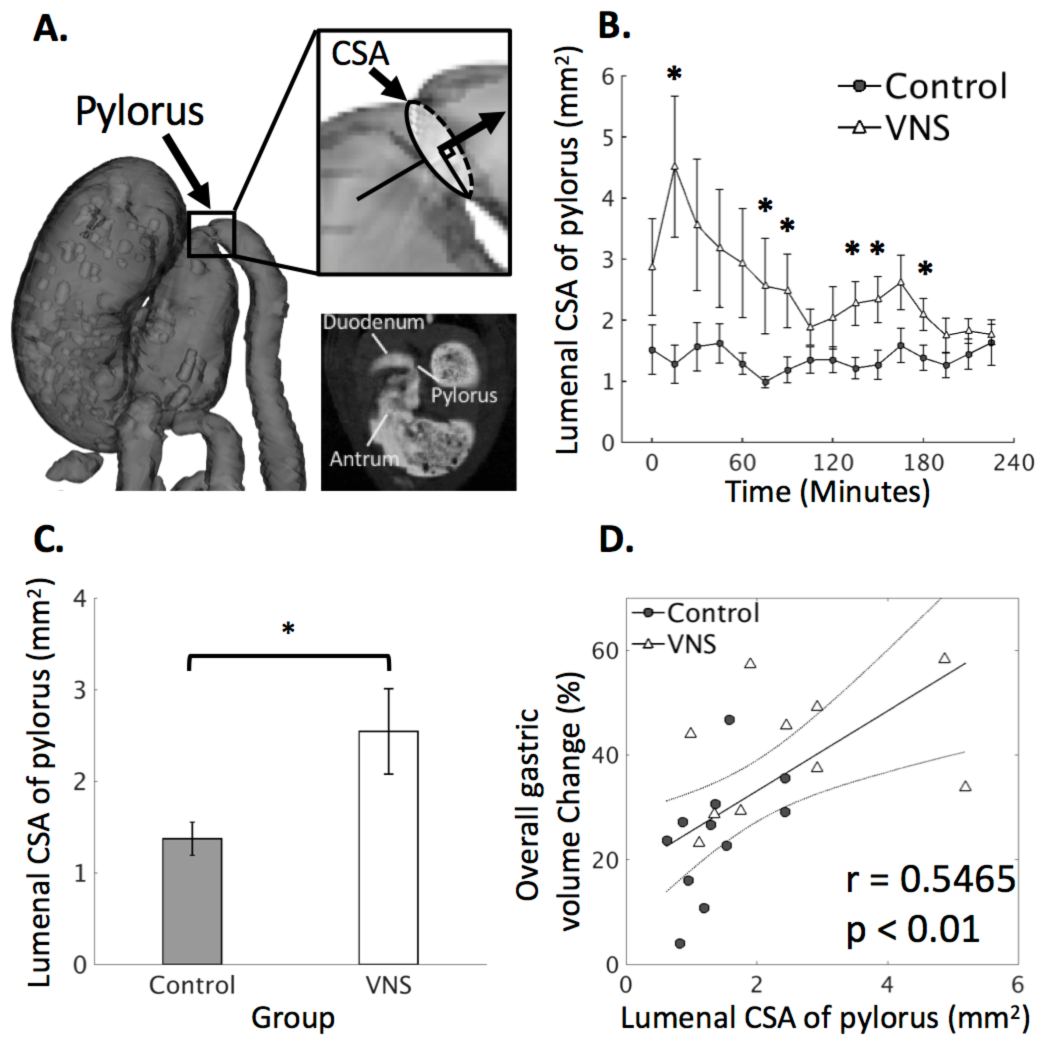
Quantification of the extent to which the pyloric ring opened under control and VNS conditions. **A.** Illustrates the location of the pylorus from a 2D slice and from a rotated 3D rendered volume. The degree to which the pyloric ring opened was measured in terms of the cross-sectional area (CSA) of the lumenal content in the pyloric canal from the high spatial resolution scans. **B.** The CSA of the pyloric ring is on average greater with VNS than without VNS, with notable and significant differences during the first 1 hour of the experiment. **C.** In general, VNS significantly increases the size of the pylorus (control vs.VNS: 1.4±0.2 vs. 2.5±0.5 mm^2^). **D.** The overall gastric volume change (%; four-hour difference) is significantly correlated to the CSA of the pyloric ring (r=0.5465, p<0.01). Noted that VNS in general leads to a larger volume change in the stomach and to a greater CSA of the pyloric ring. *: p<0.05.

### Vagus nerve stimulation did not alter antral motility

Next, we examined whether VNS modulated antral motility in terms of the frequency, amplitude and velocity of antral contractions. When sampled every ∼2 seconds, the contrast-enhanced gastric MRI captured the gastric motility as a wave of antral contraction propagating along the long axis of the antrum. No significant differences were found in the amplitude (control vs. VNS: 30.6±3.0 vs. 32.5±3.0 percent occlusion), velocity (control vs. VNS: 0.67±0.03 vs. 0.67±0.03 mm/s), or frequency (control vs. VNS: 6.3±0.1 vs. 6.4±0.2 cpm) of antral contractions (Fig. 4). The results suggest that the VNS parameter used in this study significantly accelerated gastric emptying without changing the frequency, amplitude, or velocity of antral contractions.

**Figure 4.**
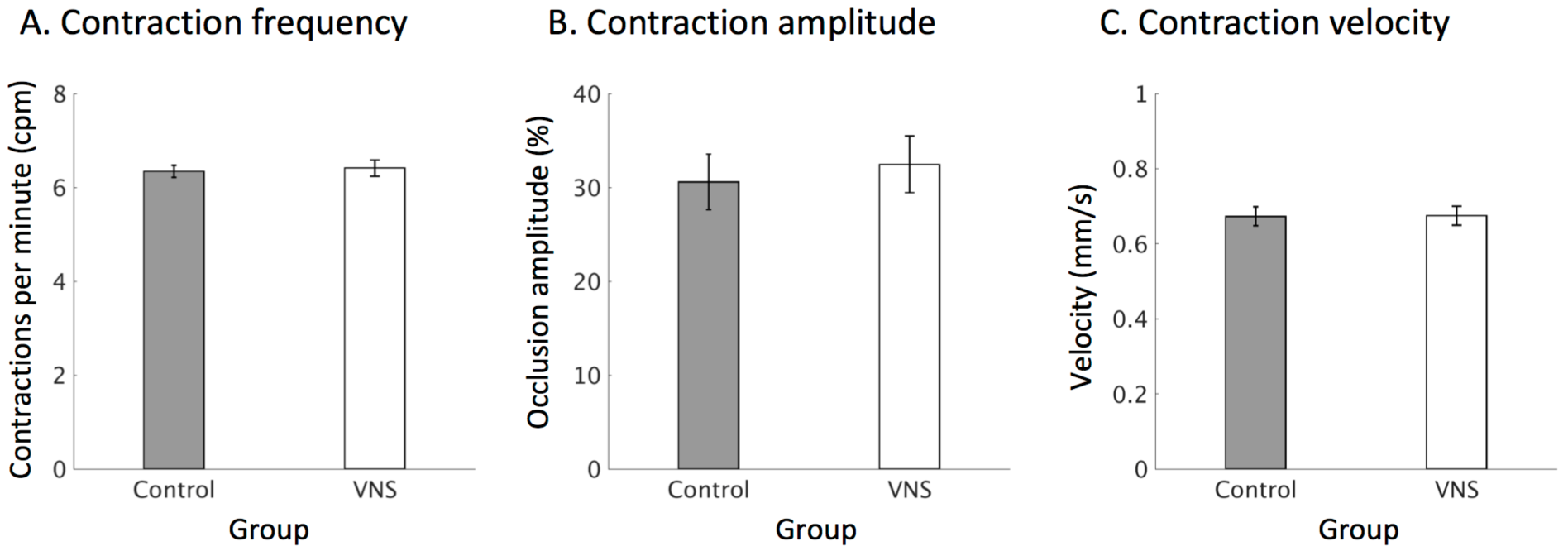
Quantification of antral motility under control and VNS conditions. A. Antral contraction frequency (control vs. VNS: 6.3±0.1 vs. 6.4±0.2 cpm). **B.** Antral contraction amplitude (control vs. VNS: 30.6±3.0 vs. 32.5±3.0 % occlusion). **C.** Velocity of peristaltic wave (control vs. VNS: 0.67±0.03 vs. 0.67±0.03 mm/s). *: p<0.05

## Discussion

Here, we used a contrast-enhanced MRI protocol to assess the effects of VNS on gastric motility and emptying in rats. We developed an effective feeding protocol in which animals voluntarily consume a Gd-labeled meal, circumventing the need of oral gavage. The Gd-labeled meal, which could be adopted for human applications, allowed gastrointestinal organs to be clearly defined. The present paradigm makes it possible to non-invasively track how the GI tract handles a voluntarily taken Gd-labeled test meal by measuring total gastric and even regional stomach emptying, antral motility (i.e., frequency, amplitude and velocity), the extent of pyloric opening, and intestinal filling.

Stimulation of the left cervical vagus nerve with the selected parameters was found to significantly enhance the rate of gastric emptying by a factor of ∼1.5. The increased emptying rate was found to primarily relate to the tonic relaxation or dilation of the pyloric sphincter, as VNS significantly enlarged the pyloric lumenal CSA (or equivalently reduced its tone). However, no significant difference was found in antral amplitude, contraction frequency, or velocity between control and VNS conditions. In sum, the experimental protocol used in this work opens an avenue for non-invasive assessment of the efficacies of neuromodulation protocols that might be used for further tuning and optimizing stimulation parameters in the future.

### Effect of vagus nerve stimulation on gastric emptying

A major finding in this study was that VNS promoted gastric emptying by increasing pyloric relaxation or opening. Two accepted driving forces pace gastric emptying. The first driving force is the transient transpyloric flow generated by antral contraction, where the peristaltic wave propagates toward the pyloric sphincter and a coordinated sequence of pyloric opening and receptive relaxation of the duodenal bulb (29, 30) leads to the forceful propulsion of chyme through the pyloric opening. The second driving force is from the transpyloric steady flow caused by the gastroduodenal pressure gradient and maintained by the tone of the proximal stomach (31, 32). To disentangle the potential mechanisms that may underlie the VNS effect that we observed, we quantified compartmental gastric volume change, which served as an indirect index of the gastric tone in different compartments. Antral motility indices and the degree to which the pyloric ring opened (i.e. the CSA of the lumenal content in the pyloric canal) were also obtained.

Central to the regulation of the transpyloric flow is the extent to which the pylorus relaxes and opens. It has been suggested that it is the opening of the pylorus, not the intraluminal pressure per se, that dominates the continuation and cessation of flow (33). Results in this study show that the chosen VNS parameter (0.6mA, 0.36ms, 10Hz) increased the degree to which the pyloric ring opened, therefore leading to a potential increase in transpyloric flow. The extent of relaxation of the pylorus was further found to correlate with gastric emptying rate. With unprecedented spatial and temporal resolution, these present MRI observations are consistent with earlier studies showing, in broad outline, that stimulation of efferent vagal fibers decreases pyloric resistance and thus increases transpyloric flow in cat (34, 35), dog (36) and pig (37) models. However, the elicited response appears to depend critically on the stimulation parameters. It is well accepted that efferent vagal fibers carry both excitatory and inhibitory stimuli to the GI smooth muscles including the pylorus. When stimulating at comparatively low threshold parameters, the excitatory (cholinergic) pathway could be selectively activated to upregulate the tone of the pyloric sphincter. At higher threshold stimulation parameters, activation of the inhibitory (non-adrenergic non-cholinergic, NANC) pathway – which in turn relaxes the pylorus (12, 22, 38) – can be superimposed on the excitatory effect of the cholinergic pathway. Martinson et al. showed that short stimulation pulse width induced excitatory effects on GI motility, while increasing the duration of the stimulatory pulses to higher values elicited inhibitory effects as well (39). Moreover, Allescher et al. demonstrated that stimulation of the vagus nerve at low (0.2-0.5Hz) vs. high frequency (>0.7Hz) elicited excitatory and inhibitory pyloric response, respectively, in anesthetized dogs (40). Taken together, both the present MRI study and the prior observations with less spatial and temporal resolution suggest that the pyloric response (i.e., relaxation of the pylorus) we observed is due to the direct innervation of efferent vagal fibers to the pyloric sphincter. Since the afferent vagal pathways were not blocked, we cannot rule out the possibility that afferent stimulation of vagus may be differentially processed in the brainstem, thus recruiting different subpopulation of efferent fibers (i.e., either excitatory, inhibitory, or both) to evoke the appropriate pyloric response.

Based on our findings and previously established factors that govern the pyloric sphincter and gastric emptying, we evaluated whether VNS elicited any excitatory or inhibitory effects on antral motility. Our results showed that VNS does not significantly alter antral motility, with perhaps only a slight increase in antral contraction amplitude. It is not unexpected to see the contraction frequency and the velocity of the peristaltic wave remain unchanged, as it is generally believed that the intrinsic gastric pace-setter potential (associated with the interstitial cells of Cajal network) sets the frequency and the velocity of the contractions in mono-gastric animals, whereas the level of vagal discharge only influences the amplitude of the contractions (41). The seemingly unchanged contraction amplitude in response to VNS may also be attributed to the specific pattern of stimulation employed. Grundy et al. showed that efferent stimulation with a constant frequency (i.e., the same pattern as we used in this study) evoked an increased tone in the ferret antrum without a notable change in the contraction amplitude (23), a result consistent with earlier findings in cat (22). However, stimulating in bursts (10 times faster for one-tenth of the time as the constant frequency stimulation group) elicited significantly greater antral contractions compared to the prestimulation condition (23). Speculatively, the main difference between these two types of stimulation is that when stimulating at continuous frequency, both excitatory and inhibitory pathway are activated, and the latter overshadows the former effect. However, when stimulating at bursts, successive post-synaptic events are likely superimposed due to temporal summation of the fast-delivered electrical pulses, therefore resulting in a dominating acetylcholine output within the enteric plexus that could elicit excitatory antral contractions. Additional investigations will be required to disentangle the effect of VNS on tonic change of the gastric antrum.

### Naturalistic feeding protocol and non-invasive imaging ensures undisturbed vagovagal reflex

A noteworthy feature of this study is that the animals were trained to voluntarily consume the test meal. Naturalistic food ingestion allows for interaction between meal and oral mucosa to properly activate cognitive and sensory processing in the brain (42, 43). These processes presumably take place in the brainstem as well as higher brain areas (e.g., amygdala), thereby supplying synaptic input to modulate the DMV neurons that control the vagal outflow to the gut. Further, meal-associated, swallow-induced esophageal distention activates vagovagal reflexes, thus triggering receptive relaxation of the forestomach and the corpus. Receptive relaxation allows the expansion of the gastric reservoir to accommodate the ingesta with minimal change of intra-gastric pressure (44). In contrast, oral gavage typically introduces stress (45, 46). In addition, artificial delivery (i.e., oral gavage) of the food directly to the stomach bypasses vagovagal reflexes and increases intra-gastric pressure (47).

This study highlights the capability of animal gastric MRI in assessing physiological functions of the stomach. In-vivo assessment of natural GI functions in small animals is more challenging given their smaller body size, much faster gastric motility, and the need for voluntary meal consumption. Existing methods are often invasive (or even lethal) (48), radioactive (49) and indirect (50), or employ other imaging methods (51) that offer limited spatial, temporal, or quantitative resolution. For example, the gastric barostat method requires intubation of a balloon in the proximal or distal stomach to measure gastric motility. Such invasive intubation in fact acts like a bolus, which could be physiologically confounding and greatly interrupts vagal innervation to the gut. Another example is ultrasound, which allows for non-invasive imaging of the gastric reservoir (51). However, ultrasound images possess limited spatial resolution and their image quality are typically degraded by speckles. The use of ultrasound in quantifying gastric functions is technically cumbersome, and the analysis is labor intensive and often observer dependent. Lastly, radioactive imaging (i.e., gastric scintigraphy) has been mainly applied to quantify gastric volumes, but the inability of repeated measurements and the relatively low sampling rate limits its use in simultaneous assessment of contractile motility. In summary, MRI overcomes the above-mentioned challenges and provides a unique opportunity to assess the gastric response to therapeutic treatments without perturbing the ongoing and spontaneous physiology.

## Acknowledgement

This work was supported in part by NIH SPARC 1OT2TR001965 and Purdue University. The authors thank Dr. Robert Phillips for discussion in surgical procedures. Authors have no conflict of interest.

## References

1. Parkman HP, Hasler WL, Fisher RS. American Gastroenterological Association technical review on the diagnosis and treatment of gastroparesis. Gastroenterology 2004; 127: 1592–1622.

2. Patrick, A Epstein O. Review article: gastroparesis. Alimentary Pharmacology & Therapeutics 2008; 27: 724–740.

3. Woods, S Seeley R. Understanding the physiology of obesity: review of recent developments in obesity research. International Journal of Obesity & Related Metabolic Disorders 2002; 26.

4. El–Serag HB. Time trends of gastroesophageal reflux disease: a systematic review. Clinical Gastroenterology and Hepatology 2007; 5: 17–26.

5. Abell, T Bernstein VK, Cutts, T et al. Treatment of gastroparesis: a multidisciplinary clinical review. Neurogastroenterology & Motility 2006; 18: 263–283.

6. Camilleri, M Toouli, J Herrera, M et al. Intra-abdominal vagal blocking (VBLOC therapy): clinical results with a new implantable medical device. Surgery 2008; 143: 723–731.

7. Sarr MG, Billington CJ, Brancatisano, R et al. The EMPOWER study: randomized, prospective, double-blind, multicenter trial of vagal blockade to induce weight loss in morbid obesity. Obesity surgery 2012; 22: 1771–1782.

8. McCallum RW, Sarosiek, I Parkman, H et al. Gastric electrical stimulation with Enterra therapy improves symptoms of idiopathic gastroparesis. Neurogastroenterology & Motility 2013; 25: 815.

9. Andrews PL, Sanger GJ. Abdominal vagal afferent neurones: an important target for the treatment of gastrointestinal dysfunction. Current opinion in pharmacology 2002; 2: 650–656.

10. Sinclair, R Bajekal RR. Vagal nerve stimulation and reflux. Anesthesia & Analgesia 2007; 105: 884–885.

11. Travagli RA, Anselmi L. Vagal neurocircuitry and its influence on gastric motility. Nature Reviews Gastroenterology & Hepatology 2016; 13: 389–401.

12. Travagli RA, Hermann GE, Browning KN, Rogers RC. Musings on the Wanderer: What’s New in our Understanding of Vago-Vagal Reflexes?: III. Activity-dependent plasticity in vago-vagal reflexes controlling the stomach. American journal of physiology Gastrointestinal and liver physiology 2003; 284: G180.

13. Qin, C Sun, Y Chen, J Foreman RD. Gastric electrical stimulation modulates neuronal activity in nucleus tractus solitarii in rats. Autonomic Neuroscience 2005; 119: 1–8.

14. Liu, J Qiao, X Chen JZ. Vagal afferent is involved in short-pulse gastric electrical stimulation in rats. Digestive diseases and sciences 2004; 49: 729–737.

15. Prechtl JC, Powley TL. The fiber composition of the abdominal vagus of the rat. Anatomy and embryology 1990; 181: 101–115.

16. Berthoud, H Carlson, N Powley T. Topography of efferent vagal innervation of the rat gastrointestinal tract. American Journal of Physiology-Regulatory, Integrative and Comparative Physiology 1991; 260: R200–R207.

17. Ben-Menachem E. Vagus-nerve stimulation for the treatment of epilepsy. The Lancet Neurology 2002; 1: 477–482.

18. George MS, Sackeim HA, Rush AJ, et al. Vagus nerve stimulation: a new tool for brain research and therapy. Biological psychiatry 2000; 47: 287–295.

19. De Ferrari GM, Crijns HJ, Borggrefe, M et al. Chronic vagus nerve stimulation: a new and promising therapeutic approach for chronic heart failure. European heart journal 2010; 32: 847–855.

20. Malow, B Edwards, J Marzec, M Sagher, O Fromes G. Effects of vagus nerve stimulation on respiration during sleep A pilot study. Neurology 2000; 55: 1450–1454.

21. de Jonge WJ, van der Zanden EP, Bijlsma MF, et al. Stimulation of the vagus nerve attenuates macrophage activation by activating the Jak2-STAT3 signaling pathway. Nature immunology 2005; 6: 844–851.

22. Martinson J. The effect of graded stimulation of efferent vagal nerve fibres on gastric motility. Acta Physiologica 1964; 62: 256–262.

23. Grundy, D Scratcherd T. Effect of stimulation of the vagus nerve in bursts on gastric acid secretion and motility in the anaesthetized ferret. The Journal of physiology 1982; 333: 451–461.

24. Eagon, J Kelly K. Effect of electrical stimulation on gastric electrical activity, motility and emptying. Neurogastroenterology & Motility 1995; 7: 39–45.

25. Schwizer, W Maecke, H Michael F. Measurement of gastric emptying by magnetic resonance imaging in humans. Gastroenterology 1992; 103: 369–376.

26. Feinle, C Kunz, P Boesiger, P Fried, M Schwizer W. Scintigraphic validation of a magnetic resonance imaging method to study gastric emptying of a solid meal in humans. Gut 1999; 44: 106–111.

27. Lu K-H, Cao, J Oleson, S Powley TL, Liu Z. Contrast Enhanced Magnetic Resonance Imaging of Gastric Emptying and Motility in Rats. IEEE Transactions on Biomedical Engineering 2017.

28. Locatelli, I Mrhar, A Bogataj M. Gastric emptying of pellets under fasting conditions: a mathematical model. Pharmaceutical research 2009; 26: 1607–1617.

29. Gleysteen JJ, Gohlke EG. The antrum can control gastric emptying of liquid meals. Journal of Surgical Research 1979; 26: 381–391.

30. Stemper T. Gastric emptying and its relationship to antral contractile activity. Gastroenterology 1975; 69: 649–653.

31. Strunz UT, Grossman MI. Effect of intragastric pressure on gastric emptying and secretion. American Journal of Physiology-Endocrinology And Metabolism 1978; 235: E552.

32. Kelly KA. Gastric emptying of liquids and solids: roles of proximal and distal stomach. American Journal of Physiology-Gastrointestinal and Liver Physiology 1980; 239: G71–G76.

33. Ehrlein H. Inductographic studies on the pyloric sphincter. Communications in Soil Science and Plant Analysis 1992; 1008: 493–493.

34. Edin, R Ahlman, H Kewenter J. The vagal control of the feline pyloric sphincter. Acta Physiologica 1979; 107: 169–174.

35. Edin R. The vagal control of the pyloric motor function: a physiological and immunohistochemical study in cat and man. Acta physiologica Scandinavica Supplementum 1980; 485: 1–30.

36. Mir SS, Telford GL, Mason GR, Ormsbee HS. Noncholinergic nonadrenergic inhibitory innervation of the canine pylorus. Gastroenterology 1979; 76: 1443–1448.

37. Malbert, C Mathis, C Laplace J. Vagal control of pyloric resistance. American Journal of Physiology-Gastrointestinal and Liver Physiology 1995; 269: G558–G569.

38. Martinson J. Vagal Relaxation of the Stomach Experimental Re-investigation of the Concept of the Transmission Mechanism. Acta Physiologica 1965; 64: 453–462.

39. Martinson, J Muren A. Excitatory and inhibitory effects of vagus stimulation on gastric motility in the cat. Acta Physiologica 1963; 57: 309–316.

40. Allescher, H Daniel, E Dent, J Fox, J Kostolanska F. Extrinsic and intrinsic neural control of pyloric sphincter pressure in the dog. The Journal of physiology 1988; 401: 17–38.

41. Szurszewski JH. Modulation of smooth muscle by nervous activity: a review and a hypothesis. Federation proceedings 1977: 2456–2461.

42. Bonnichsen, M Dragsted, N Hansen AK. The welfare impact of gavaging laboratory rats. Animal Welfare-Potters Bar Then Wheathampstead- 2005; 14: 223.

43. Brown AP, Dinger, N Levine BS. Stress produced by gavage administration in the rat. Journal of the American Association for Laboratory Animal Science 2000; 39: 17–21.

44. Rogers, R Hermann, G Travagli R. Brainstem pathways responsible for oesophageal control of gastric motility and tone in the rat. The Journal of Physiology 1999; 514: 369–383.

45. Stengel, A Taché Y. Neuroendocrine control of the gut during stress: corticotropin-releasing factor signaling pathways in the spotlight. Annual review of physiology 2009; 71: 219–239.

46. Zheng, J Dobner, A Babygirija, R Ludwig, K Takahashi T. Effects of repeated restraint stress on gastric motility in rats. American Journal of Physiology-Regulatory, Integrative and Comparative Physiology 2009; 296: R1358–R1365.

47. Damsch, S Eichenbaum, G Tonelli, A et al. Gavage-related reflux in rats: identification, pathogenesis, and toxicological implications. Toxicologic pathology 2011; 39: 348–360.

48. Uchida, M Iwamoto C. Influence of amino acids on gastric adaptive relaxation (accommodation) in rats as evaluated with a barostat. Journal of Smooth Muscle Research 2016; 52: 56–65.

49. Jordi, J Verrey, F Lutz TA. Simultaneous assessment of gastric emptying and secretion in rats by a novel computed tomography-based method. American Journal of Physiology-Gastrointestinal and Liver Physiology 2014; 306: G173–G182.

50. Uchida, M Kobayashi, O Saito C. Correlation Between Gastric Emptying and Gastric Adaptive Relaxation Influenced by Amino Acids. Journal of neurogastroenterology and motility 2017; 23: 400.

51. Gilja, O Hausken, T Wilhelmsen, I Berstad A. Impaired accommodation of proximal stomach to a meal in functional dyspepsia. Digestive diseases and sciences 1996; 41: 689–696.

